# The *Incubascope* : a simple, compact and large field of view microscope for long-term imaging inside an incubator

**DOI:** 10.1101/2021.09.21.461183

**Authors:** Amaury Badon, Laetitia Andrique, Amaël Mombereau, Louis Rivet, Adeline Boyreau, Pierre Nassoy, Gaëlle Recher

**Author notes:** **optical microscopy**.

## Abstract

Optical imaging has rapidly evolved in the last decades. Sophisticated microscopes allowing optical sectioning for 3D imaging or sub-diffraction resolution are available. Due to price and maintenance issues, these microscopes are often shared between users in facilities. Consequently, long term access is often prohibited and does not allow to monitor slowly evolving biological systems or to validate new models like organoids. Preliminary coarse long-term data that do not require acquisition of terabytes of high-resolution images are important as a first step. By contrast with expansive all-in-one commercialized stations, standard microscopes equipped with incubator stages offer a more cost-effective solution despite imperfect long run atmosphere and temperature control.

Here, we present the *Incubascope*, a custom-made compact microscope that fits into a table-top incubator. It is cheap and simple to implement, user-friendly and yet provides high imaging performances. The system has a field of view of 5.5 × 8 mm^2^, a 3 *μ*m resolution, a 10 frames/second acquisition rate, and is controlled with a Python-based graphical interface. We exemplify the capabilities of the *Incubascope* on biological applications such as the hatching of *Artemia salina* eggs, the growth of the slime mold *Physarum polycephalum* and of encapsulated spheroids of mammalian cells.

## Introduction

Optical microscopy plays a central role in modern biology because it allows to image samples with low photo-damage and at high spatial and temporal resolution (1). It provides a unique opportunity to observe sub-cellular structures or processes over scales that span millimeters or centimeters. In the last few years, numerous commercial systems have been developed to image living specimens with higher resolution, deeper into the tissues, with or without exogenous and specific contrasts. Yet, optical imaging of living samples remains challenging for several practical reasons. First, the phototoxicity related to fluorescence emission may become significant in long term acquisitions (2). Second, and often overlooked, the necessity to perform imaging in environmental conditions that are as close as possible from the physiological conditions (temperature, humidity, carbon dioxide and/or oxygen partial pressure) is critical when one deals with mammalian cells that require sterile conditions and whose behaviour is strongly affected by the surrounding atmosphere and temperature. The most widely used method is to enclose the microscope in a commercially available incubator box (23–25). This option is satisfactory for image acquisition no longer than typically 2 days because it does not offer optimal gases and temperature control and sterility maintenance as they could be reached in a culture room incubator. As a corollary, for longer term imaging (from days to weeks), besides the above mentioned difficulties, an additional drawback is that the microscopes are immobilised for the time course of the experiment, which is often prohibitive for cost and management reasons in imaging facilities. The solution may consist in keeping the sample in a standard incubator and moving it back and forth to the microscope equipped with an incubator box every hour or day. This alternative is not only time-consuming but is also not well-suited for multi-positions imaging since the exact same location on the specimen has to be searched at each time point.

Academic researchers and microscopy companies sought to develop compact boxes comprising both the microscope module and the control of the environmental conditions or to fit directly imaging devices inside an incubator. Compact hybrid platforms are often dedicated to high-content screening studies. Numerous companies have developed such sophisticated and expensive “one-button solution” apparatuses (20–22) and target big pharma companies or facilities. Other all-in-one cell analysis systems (3, 19, 27)) are more affordable for academic laboratories. Some very cost-effective approaches offer relatively high optical performances but with specific limitations. For instance, lens-free standalone systems may require numerical reconstructions that are sometimes artefactual (31), while other systems working only in transmission mode are not best suited for imaging thick tissues.

In this context, there is a need for a system that gathers the following requirements: compact design, hybrid (fluorescence and bright-field) acquisition, amenability to high-content and ease of use for biologists.

Practically, we developed an imaging system that fits within a bench-top incubator. It offers a very large field of view (FOV = 5.5 × 8 mm^2^) with a sub-cellular spatial resolution (∼ 3 *μ*m). Our system is cost-effective thanks to a frugal architecture made of a few commercially available optical and electronic elements and an open source acquisition code. Two imaging modalities, bright-field and epifluorescence, provide complementary information on the sample of interest. In this work, we report a detailed description of our system, coined the *Incubascope*, in order to allow replication or adaptation by other research groups. We exemplify the performances of the *Incubascope* by imaging several very different biological systems. Using wild and self-sustainable small crustacean eggs (*Artemia salina*), we show that we could monitor the hatching process and swimming motions of the larva. Then, using the plasmodial slime mold *Physarum polycephalum*, better known as blob, whose growth exhibits multi-scale branching patterns, we illustrate the advantage of the *Incubascope* to perform simultaneous epifluorescence and bright-field microscopy over a large field of view. Finally, we imaged aggregates of eukaryotic cells or multicellular spheroids encapsulated within alginate capsules that can serve as tumor model mimics (7). Since mammalian cells typically divide once every day, growth monitoring requires imaging over several days in optimal physiological conditions. Altogether, the *Incubascope* as designed and validated in the present work is expected to face some unmet needs in biological imaging.

## Materials and Methods

In this section, we describe the design and working principle of the *Incubascope*. We detail (a) every component of the proposed system and its assembly, (b) the open-source software used to control the acquisition parameters, (c) the optical performances of the apparatus, (d) the biological samples used for experimental validation and (e) finally the 3D printed parts used to accommodate the samples.

### Parts and components

The *Incubascope* has two imaging modes, namely bright-field and epifluorescence. In the configuration presented h ere, the fluorescence excitation wavelength is centred around 490 nm and the wavelength of the transmitted light is centred around 625 nm. Other configurations can be obtained by simply changing the illumination sources and filters depending on the specificities of the samples and probes.

#### Opto-mechanical components

Fluorescent excitation is provided by a single mounted LED (M490L4, Thorlabs) (Figure 1, LED 1), collimated with an aspheric lens (L2) and reflected by a 505 nm longpass dichroic mirror (DMLP505R, Thorlabs). Light is focused at the back aperture of a low magnification microscope objective (TL2X-SAP, Thorlabs). This objective combines a very large field of view (FOV=11 mm), a large working distance (56 mm) and a relatively high numerical aperture (NA=0.1) thus providing a lateral resolution up to 2.5 *μ*m. As the large working distance can be problematic to keep a compact system, we decided to install the MO horizontally with a mirror at 45° in between the MO and the sample. In this geometry, the physical height of the system is reduced and the whole apparatus can sit on a single rack in a table-top incubator (see Figure 1.A and B). The sample mount consists in a lens mount (CP33, Thorlabs) attached on a 1 axis stage to adjust manually the focus. Light for the bright-field (or transmission) mode is provided by a second mounted LED (M625L4, Thorlabs) (Figure 1.D, E LED2), collimated with an aspheric lens (L1) and reflected by a mirror placed at 45° above the sample.

**Fig. 1.**
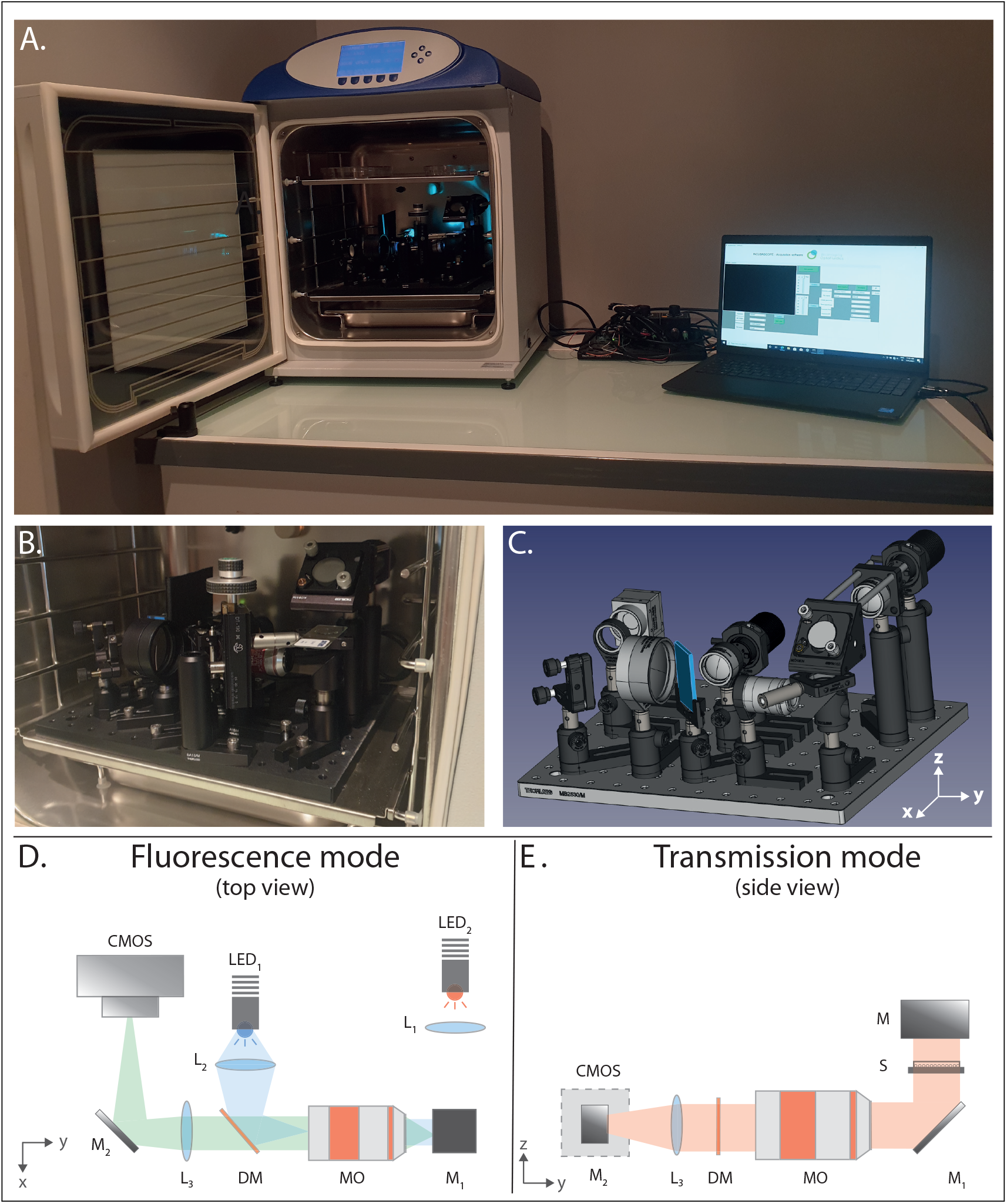
Optical layout of the *Incubascope*. (a) Photography of the prototype placed on a single rack inside a table-top incubator. The *Incubascope* is controlled by a laptop using only USB cables and an Arduino board. (b) Photography of the *Incubascope* showing the optical components placed on a single breadboard. (c) 3D rendering of the optomechanical components placed on a single breadboard inside the incubator. (d-e) Schematic of the apparatus in fluorescence and transmission modes respectively. DM : dichroic mirror, LED : light emitting diode, L : lens, M : mirror, MO : microscope objective, S : sample.

For both modes, the light is collected by the MO, transmitted through the DM and focused by an achromatic doublet (L3) that acts as a tube lens (AC508-150-A-ML, Thorlabs). A mirror at 45° is inserted between this lens and the camera to fold the beam path and spare space.

#### Electronic components

In order to simultaneously reduce the cost of the system and ensure a long viability inside the incubator, we decided to reduce the number of electronic components placed in the incubator. Hence, our system, which is primarily dedicated at performing large field of view imaging at low to moderate magnification, does not include actuators to control the position of the sample laterally or axially. It also reduces the risk of misalignment.

Inside the incubator, a CMOS camera is placed at the imaging plane of the tube lens (L3). This camera needs to be small, USB connected, cost-effective and with a large number of pixels. Indeed, the MO used in our system offers a very large FOV with a moderate spatial resolution. The amount of spatial information transmitted by this MO can be quantified by its spatial bandwidth product (SBP) (14–16). This quantity defines the number of pixels required to captured the whole amount of information with a Nyquist sampling. In practice, we obtain :

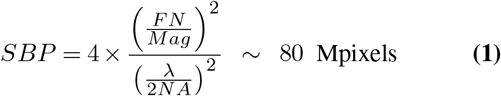

with FN the field number of the MO and Mag its magnification. In theory, a 80 Mpixels sensor is necessary to collect all the information captured by the MO for a 500 nm wavelength. Yet, cameras with this amount of pixels are very expensive (> 20 k €), not USB connected and bulky. A cost-effective 20 Mpixels (Basler acA5472-17uc, see Table 1) was chosen for its number of pixels/price ratio.

**Table 1.** Detailed list of the components of the *Incubascope* and their prices.

| Name | Component | Product code | Supplier | Quantity | Cost per unit (€) |
| --- | --- | --- | --- | --- | --- |
| CMOS | USB monochrome camera | acA5472-17um | Basler | 1 | 550 |
| ARD | Arduino / DAQ board | Arduino Uno | Arduino | 1 | 20 |
| LED 1 | 490 nm mounted LED | M490L4 | Thorlabs | 1 | 200 |
| LED 2 | 625 nm mounted LED | M625L4 | Thorlabs | 1 | 200 |
|  | LED drivers | LEDD1B | Thorlabs | 2 | 294 |
| MO | Microscope objective | TL2X-SAP | Thorlabs | 1 | 1100 |
| TL | Tube lens | AC508-150-A-ML | Thorlabs | 1 | 135 |
| DM | dichroic mirror | DMLP505R | Thorlabs | 1 | 230 |
| FL | fluorescence filter | MF525-39 | Thorlabs | 1 | 100 |
|  | breadboard | MB3045/M | Thorlabs | 1 | 180 |
|  | mechanical components |  | Thorlabs | 1 | 200 |
|  |  |  |  | **Total** | **5500** |

Outside the incubator, the system includes two LED drivers (T-cube, Thorlabs) connected to an Arduino Uno board with BNC cables. The Arduino itself is connected to a computer using an USB cable which allow to remotely control the output power of both LEDs.

### Open source software for acquisition control

Preview mode and control of the acquisition parameters are performed using a custom written code in the freely available Python language. This allows to further reduce the cost of the systemand to ensure open access and possible duplication by any other research group. This was also at stake when it deals with Open-Science and making sure our device could be duplicated by anyone. Briefly, the code allows to communicate both with the camera using the Basler pypylon library (10) and with the Arduino using the Firmata library (11). Acquisition parameters such as the frame rate, exposure time,file name, repository location and data output of the camera as well as LEDs output power can be controlled using a graphical user interface. Preview live modes and long-term time lapse acquisition are provided. Our code gives the choice to save the experimental data in two different formats, tiff or hdf5 file (12). The latter is particularly interesting when dealing with large amount of data as its structure allows to open only subsets of a file. In practice, low-resolution images of a time lapse can be opened quickly to check the status of the sample while the experiment is still on. The code is publicly available in our GitHub repository (9).

### Characterization of the optical performances

The resolution and the size of the FOV of the system were first investigated using a USAF 1951 resolution test target in the bright-field mode. First, from the results displayed in figure 2, we estimate the FOV to be equal to 5.5 × 8 mm^2^. Second, the smallest resolvable line is the element 3 of the group 7. This corresponds to a spatial resolution of roughly 3.1 *μ*m. Given the magnification of the system and the pixel size, the theoretical resolution of the *Incubascope* is 3.2 *μ*m. Thus, the experimental value is in agreement with the theoretical one. From the same experiment, the collection efficiency across the FOV is investigated. From the background intensity plotted in Figure 2.C, a signal decrease of only 17% is observed over 8 millimeters which demonstrates the relative homogeneity of both the illumination and the collection efficiency of the system.

**Fig. 2.**
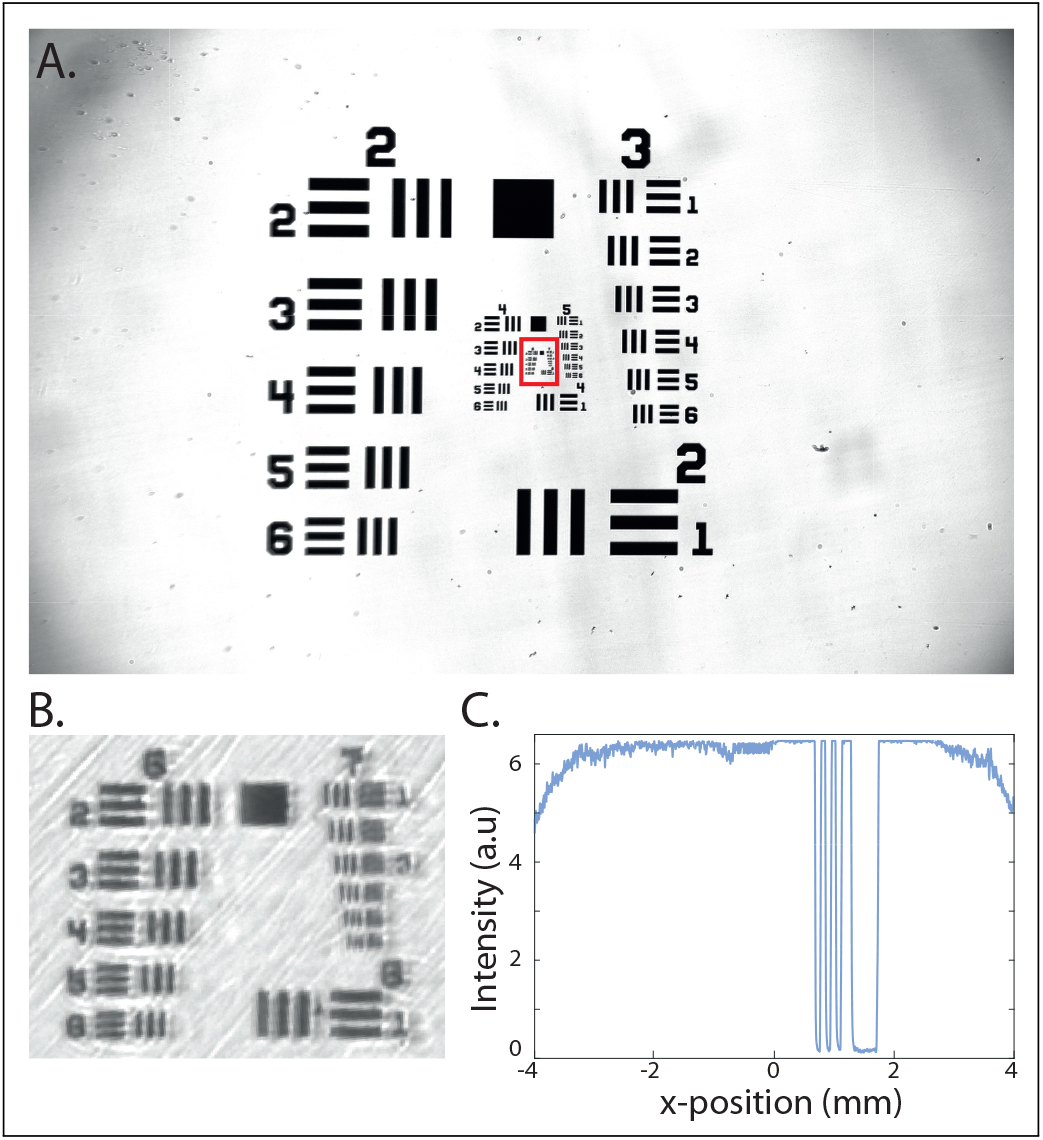
Optical performances of the *Incubascope*. (a) Bright-field i mage o f a USAF resolution target. (b) Zoom in of the area highlighted by the red square in (a). The smallest resolvable line is the element 3 of the group 7. (c) Plot of the intensity profile along a line of the image displayed in (a). A decrease of 17 % is observed across the 8 mm FOV.

Additionally, the mechanical stability of the *Incubascope* over time was evaluated similarly to the procedure described in (8). The signal corresponding to the corner pixel of the opaque square on the test target was evaluated over time. As seen on figure S 1, n o m echanical d rift w as o bserved over hours both laterally and axially. Both the high mechanical performances of the system adn the large DoF of the microscope objective may explain this remarkable stability.

### Biological samples

#### Artemia eggs

Artemia (*Artemia salina*) eggs were purchased as a powder (Jeulin, Evreux, France). They were revitalised in salted water (30 g.L^−1^), according to provider’s recommendations.

#### Blob

Blobs (*Physarum polycephalum*) were purchased as dried sclerotium (Blobshop, Toulouse, France). They were passaged on agar-agar plates and fed with dried oat flakes (BioCoop, Talence, France). Fluorescence labelling was achieved after dropping 200 *μ*L of a Bodipy solution (D3921, ThermoFisher) on the blob, allowing for 30 min incubation and washing with running water.

#### Cellular Capsules

We used the HEK-293T (gift from A. Bikfalvi lab) cell line that is obtained from embryonic kidneys and known to be highly proliferative. Cells were infected with lentiviruses to stably and ubiquitously express cytoplasmic GFP labelling. Hollow alginate capsules were produced as previously described (7). Briefly, a co-extrusion three-way microfluidic chip was used to generate composite droplets in the air, which readily crosslink when entering into a 100 mM CaCl_2_ gelation bath and encapsulate the cell suspension in a nutrient-permeable shell. Cellular capsules were grown in regular DMEM Glutamax medium (10% SVF and 1% Penicillin/Streptomycin).

### 3D printed parts

A homemade agarose-based basin was fabricated for the *A*. *salina* eggs experiments to ensure that the hatched shrimps remain in the FOV. In a 35 mm petri dish, 2% agarose solution was imprinted with a stamp, which was drawn with SolidWorks (Dassault System). The main pool (3 mm-depth) is designed so that a thin layer of agarose remains at the bottom, preventing any leak. The stamp was printed with a DLP stereolithography printer (MicroPlus HD printer, EnvisionTec, HTM-140 resin). Another stamp was used for the spheroids experiments to order the HEK-filled capsules and allow individual monitoring. The fabrication principle of the stamp is the same as for the basin one, and the shape of individual microwells was designed as described in (17). Both STL files are provided as Supplemental materials.

## Results

### Monitoring egg hatching and swimming activity of *A salina*

As a first experimental validation of the *Incubascope*, we observed the hatching of population of *A salina* eggs. The organisms first hatch within hours to a few days, before they start swimming. Thus, long but also fast rate acquisition is necessary to monitor their birth and trajectories.

Around 60 dried eggs were deposited in the dedicated basin (see figure 3.A) and observed for 2 days. These specimens are not fluorescently labelled. Only bright-field images with an exposure time of 20 *μ*s were taken with various time intervals.

**Fig. 3.**
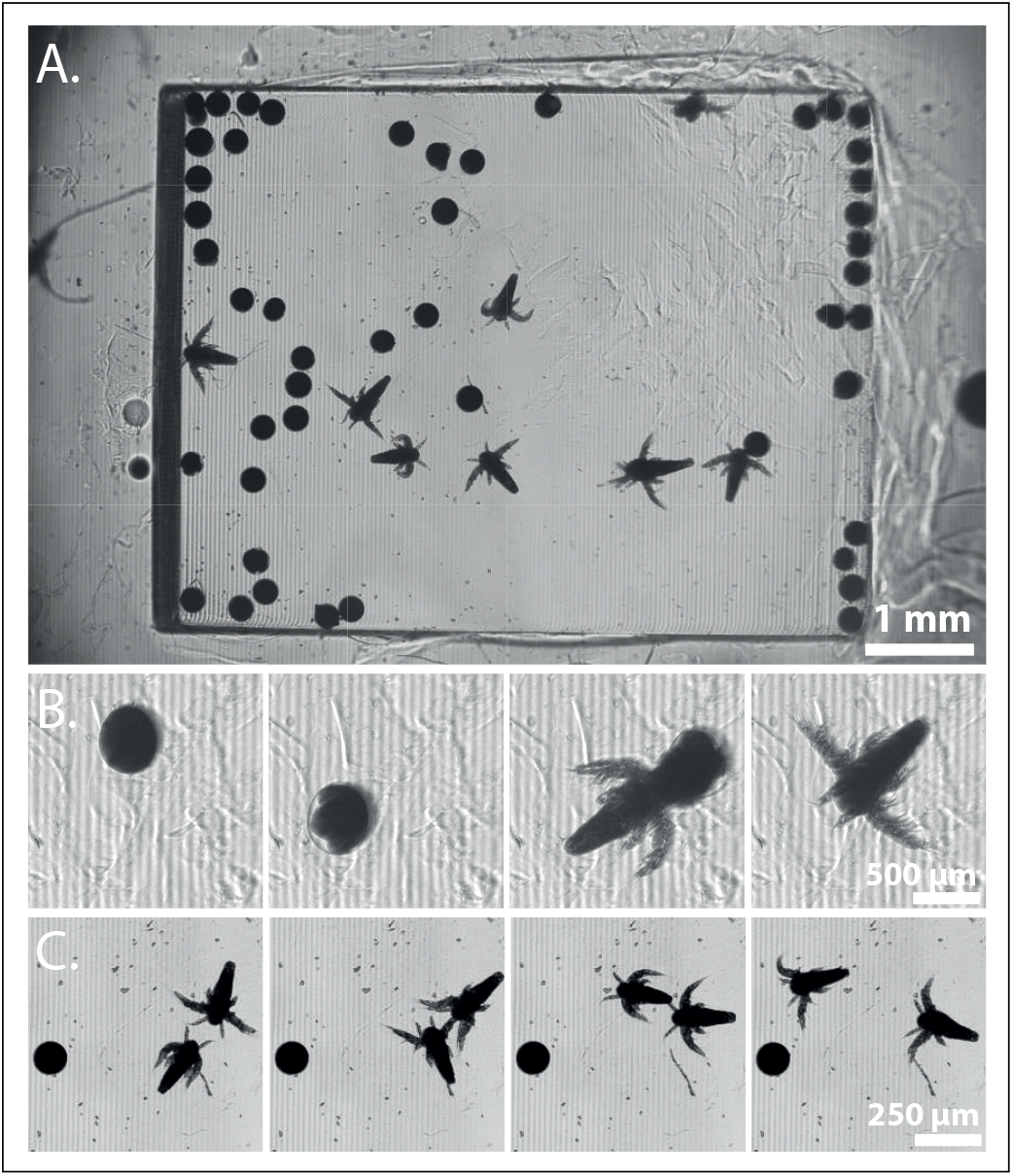
Observation of a population of *A*. *salina*. (a) Bright-field image showing the large cuvette filled with eggs and young *A*. *salina*. (b) Time lapse displaying the hatching of an egg. Images are acquired every minute. (c) Time lapse displaying two specimens of *Artemia* swimming in opposite directions and collided. Images are acquired every 100 ms.

First, we extracted frames acquired every minute to show the hatching of one egg. As seen on figure 3.B, both the temporal and the spatial resolutions enable to follow this event. Note that eggs and hatched shrimps move in 3D in this deep swimming pool. This is the reason why, even though our system has a large DoF, an egg can be in focus at a given frame but slightly blurred at the next one.

Then, we captured images at the highest frame rate possible with our system. In theory the camera can acquire images up to 17 frames per second (fps), but this value is reduced in practice to 10 fps with our acquisition software. These performances are however sufficient to precisely capture the displacements of mature *A*. *salina* across the basin. Figure 3.C displays four consecutive frames showing a collision between two specimens swimming in opposite directions. More data are provided in the supplementary information.

### Imaging the migration of *Physarum polycephalum*

In the second explored case study, the imaging challenge is no longer to follow a population of individuals but the development of an extended specimen. The blob (*P*. *polycephalum*) was chosen because it exhibits a multiscale and dynamic cyto-architecture (18, 28–30).

To be able to monitor a whole migration process, which occurs through the formation of a large and dynamic network of vessels, we started with a rather tiny blob, and we placed an oat flake a few millimetres away in the camera FOV. We labelled the blob with a lipophilic fluorescent d ye (Bodipy 505/515) to monitor the reshaping of the vessels in the fluorescence channel. As shown in figure 4, the vessels are well delineated and the lumen appears as a dark strip flaked with bright walls.

**Fig. 4.**
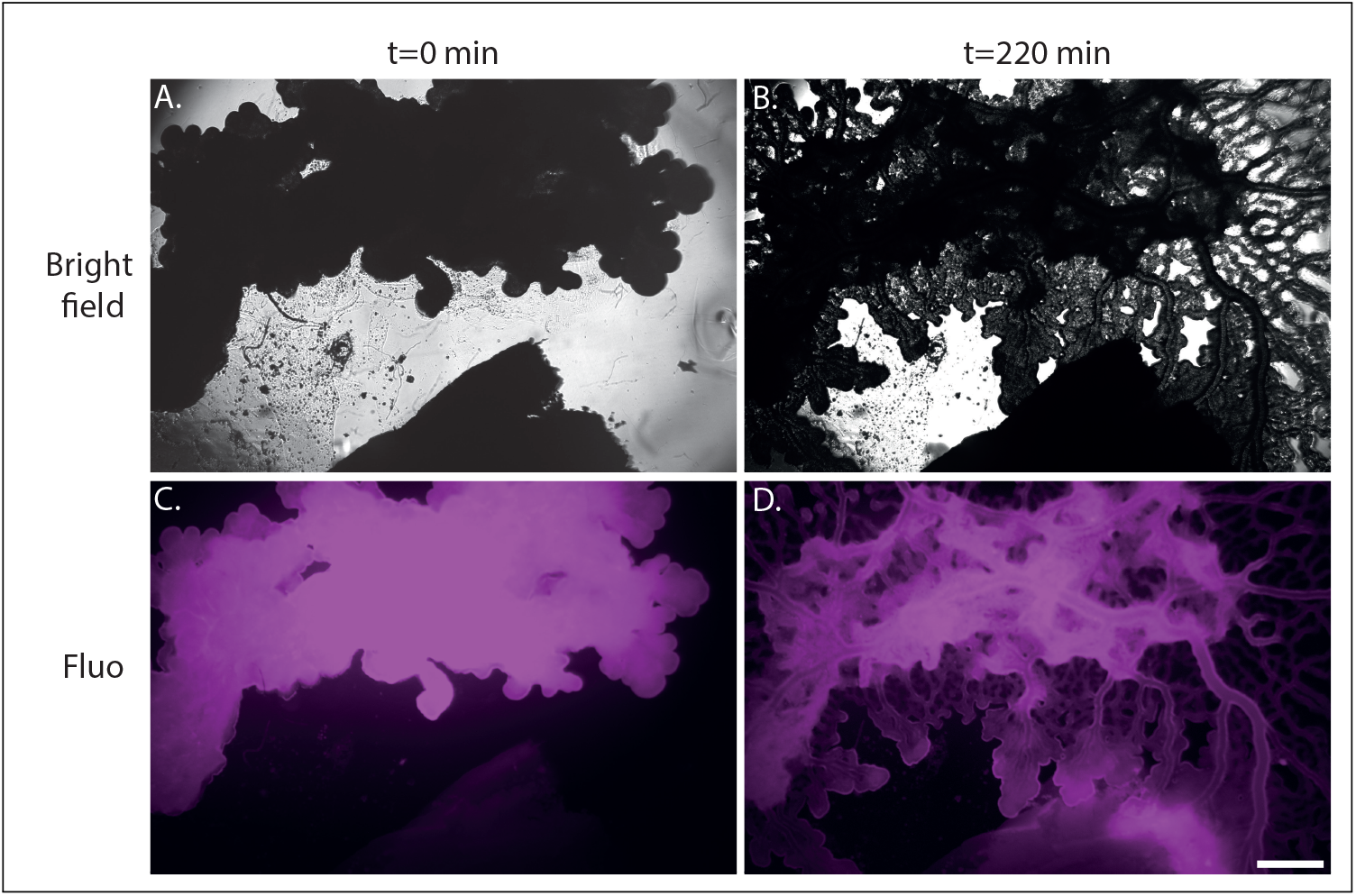
Long-term observation of *P*. *polycephalum*. (a,c) Bright-field and fluorescence images of a *P*. *polycephalum* specimen (on top) moving towards an oat flake (on the bottom) at t=0h and for t=220 min (b,d).

### Imaging a large population of multicellular spheroids

Third, we used 3D cell cultures of GFP labelled HEK-293T cells. More specifically, we prepared multicellular spheroids encapsulated in individual hydrogel shells as described in (7) and summarized in the Materials and Methods section. Since mammalian cell doubling time is of the order of a day, imaging over several days is required to investigate their growth. Transmission and epifluorescence images were acquired every hour during 72 hours with exposure times equal to 22 *μ*s and 500 ms respectively.

As seen in figure 5.A and B, hundreds of spheroids of about 300 *μ*m in diameter were imaged within the FOV. Due to the large number of pixels of the CMOS sensor, a close up view on a single spheroid provides a precise visualization of the sample (see figure 5.C,D). Cell proliferation leading to the increase in size of the spheroids inside the alginate capsule could be monitored at least for 3 days. The large number of spheroids simultaneously observed within the same field of view allows to compare their growth in the same conditions but also to detect isolated behaviours. For instance, the rupture of a capsule can be observed when cells overfill the core of the capsule (see supplementary information) or/and when a thinning in the alginate wall represents a weak-point (Figure 5.D).

**Fig. 5.**
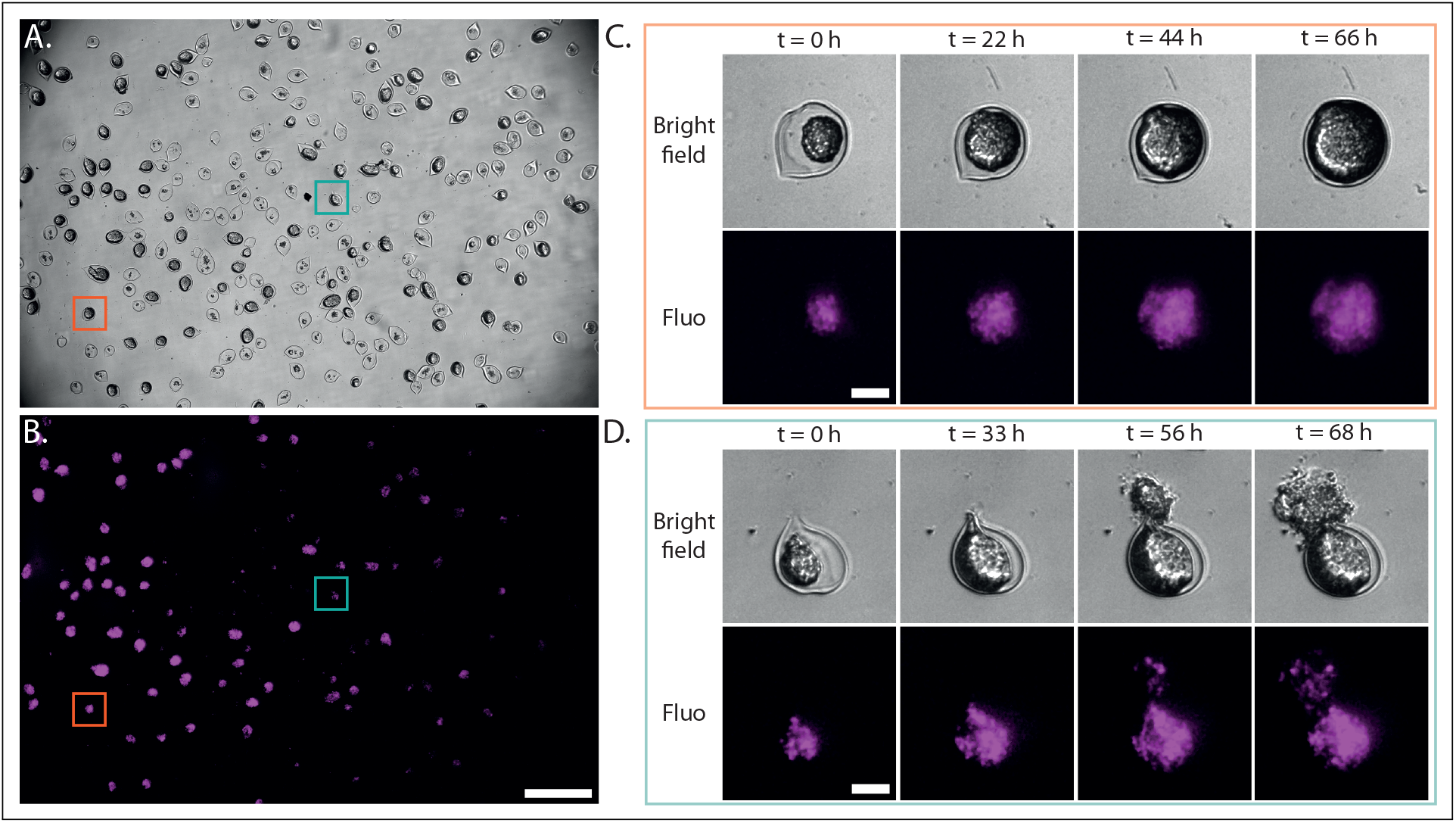
Long-term observation of spheroids. (a) Bright-field image showing hundreds of spheroids placed in a Petri dish. (b) Same as (a) but in fluorescence mode. (c) Time lapse in both modes of a single spheroid highlighted by the orange square in (a). Cell proliferation leads to capsule overfilling. (d) Time lapse in both modes of a single spheroid highlighted by the green square in (a). Here, a defect in the capsule wall induces the bursting of the shell when the pressure becomes to high.

## Discussion

In the present work, we have first provided a detailed description of the components, the open-source acquisition software, the optical performances of the *Incubascope*. Then, we have chosen three different experimental systems to exemplify the versatility and performances of the *Incubascope*. Our custom-built apparatus offers a simple, compact and cost-effective solution for long-term observations of biological samples inside a table-top incubator. With a 5.5 × 8 mm^2^ FOV and a 3*μ*m spatial resolution, our system is a two-in-one device and an ideal imaging system for the observation of a diversity of biological samples.

In this section, we now want to discuss the cost of the system, the potential solutions to further reduce it as well as the current imitations.

### Costs

Table 1 shows a detailed list of the items and costs. The cost of the entire system is approximately 5500 C and includes the optics, the electronics, the light sources and the mechanical parts, but excludes the price of the incubator. The code used to control and perform acquisitions is written in the open source Python language, and thus does not require to have access to a specialized software licence (e.g Matlab or Labview).

Light source components and associated drivers cost around 1000 €. To further reduce the cost of the system, we may envision to use unmounted LEDs such as Adafruit light sources, which are powerful, compact and easily controlled with an Arduino board.

Regarding the optical elements, cost-effective solutions had been proposed to replace the MO by a single lens or even to use lens from cell phones. Yet these solutions usually diminish the optical performances and also require to adapt components or dismantle commercial products. By contrast, our approach was to solely use commercially available optical elements for the *Incubascope*.

Finally, one could also print all the optomechanical parts with a high-quality 3D-printer (26). This inexpensive way to build a prototype is *a priori* attractive, although mechanical stability will have to be carefully tested.

### Limitations and potential improvements

Our system was initially developed to image the dynamics of a large number of 3D cell cultures during several days. In this context, we chose to maximize the FOV at the cost of a moderate - but sub-cellular - resolution. For other applications a higher magnification MO would be required. Yet, our current configuration is designed for a MO with a 95 mm parfocal length and a focal length large enough to insert a 45° mirror between the objective front lens and the sample. Hence, adapting our system for other objectives will require major modifications of the layout.

Secondly, we presented imaging results with both fluorescence and bright-field modes. In the current configuration, we decided to excite the fluorescence with a 490 nm LED well suited to our own needs Note that multiple fluorescent signals can be obtained simultaneously by adding more LEDs and coupling them using dichroic mirrors or beam splitters in the illumination path as done in (8) for instance, and by replacing the current long-pass dichroic mirror by a multi-band one. Yet, this additional imaging modes will increase the complexity of the system, its size, but also the cost as the LED’s drivers are one of the most expensive components of the system.

In the perspective of making a cheap but robust system, we did not include any motorized stage control as the temperature and, more importantly, the humidity inside the incubator are not compatible with a long-term use of electronic components. In fact, we did not even include a manual stage to control the sample lateral positions for two reasons. First, it saves space, which is crucial for the system to fit on a single shelf of a small incubator. Secondly, as the FOV of the system is large ( ∼ 8 × 5.5 mm^2^), positioning the sample on the holder can be done manually. The axial position of the sample needs to be finely adjusted using a manual stage with a micrometer resolution. Once the focus was adjusted at the onset of the experiment, no mechanical drift that would require an auto-focus mechanism on the system was detect-ed afterwards. Here, the large depth of field of the MO ( ∼ 50 *μ*m for NA=0.1) ensured that the sample stays inside the DoF of the MO even if small axial drifts occur during an experiment.

Finally, as the acquisition software is not demanding in computer resources but only in memory disk (39 Mo/image), we plan to control the *Incubascope* with a Raspberry Pi for future works. This will further reduce the cost as well as the physical size of the system outside the incubator.

## Conclusion

The *Incubascope* presented in this work is a custom-made microscope with an very large field-of-view o f ∼ 5.5 × 8 mm^2^ and spatial resolution of ∼ 3 *μ*m that is sufficient to observe sub-cellular structures. More-over, its compact size and low cost make of the *Incubascope* a system that can be easily integrated into an incubator to perform long-term observation of the dynamics of a large ensemble of cells or 3D cell cultures. With an epifluorescence a nd a transmission mode, both structural and functional imaging can be performed over weeks. This apparatus is expected to be useful for a variety of applications that demand in parallel monitoring of numerous samples such as in tissue engineering, embryogenesis and drug discovery.

## Funding Information

This work was supported by the Agence Nationale pour le Rcherche (MecaTiss,ANR- 7-CE30-0007-03, Parkington,ANR-17- CE18-0026-02, MecanoAdipo, ANR-17-CE13-0012-01), Cancéropôle GSO, SIRIC Brio (Bordeaux) and the following charities: Association pour la Recherche sur le Cancer (ARC), La Ligue Contre le Cancer (Gironde and Pyrénées Atlantiques). A. Badon thanks the Fondation des Treilles for its financial support.

## Acknowledgments

We acknowledge Pierre Bon for fruitful discussions, our interns Margaux Caumont and Aude Gadenne, Antony Lee for help with Python, Nadège Pujol and Andreas Bikfalvi for providing HEK cells.

## Notes

### Competing Interest Statement

The authors have declared no competing interest.

