## Supplementary figures and images for "The *Incubascope* : a simple, compact and large field of view microscope for long-term imaging inside an incubator"

### Supplemental figure 1

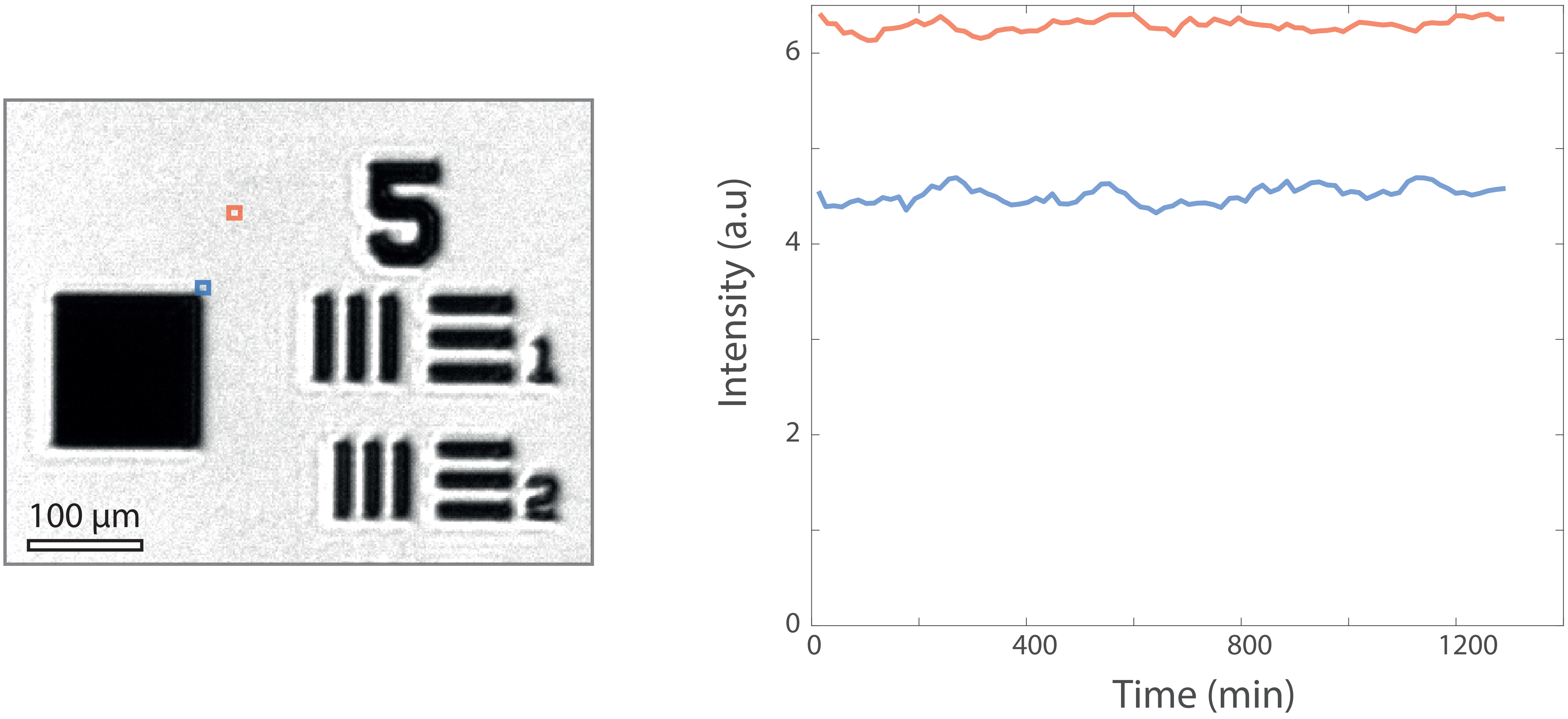
